# Global gene repression by Dicer-independent tRNA Fragments

**DOI:** 10.1101/143974

**Authors:** Canan Kuscu, Pankaj Kumar, Manjari Kiran, Zhangli Su, Asrar Malik, Anindya Dutta

## Abstract

tRNA derived RNA fragments (tRFs) is an emerging group of small RNAs as abundant as miRNAs, and yet their roles are not well understood. Here, we focus on endogenous tRFs (18-22 bases) derived from 3’ end of human mature tRNAs (tRF-3) and their functions in gene repression. tRF-3 levels increase upon parental tRNA over-expression or tRNA induction by c-Myc oncogene activation. Elevated tRF-3 levels lead to repression of target genes with a sequence complementary to the tRF-3 in the 3’ UTR. The tRF-3-mediated repression is Dicer-independent, Argonaute-dependent and the targets are recognized by 5’ seed sequence rules similar to miRNAs. Furthermore, tRF-3s associate with GW proteins in P-bodies. RNA-seq identifies the endogenous target genes of tRF-3s that are specifically repressed upon tRF-3 induction. Overall, our analysis shows Dicer-independent tRF-3s, generated upon tRNA upregulation such as c-Myc overexpression, regulate gene expression globally through Argounate via seed sequence matches.

## Introduction

There are many different classes of cellular small RNAs, including microRNAs (miRNAs), Piwi-interacting RNAs (piRNAs) and tRNA related fragments (tRFs). tRFs are 14-32 bases long RNAs derived from tRNAs that have been identified from bacteria to human with high abundance. tRFs can be classified into three groups: tRF-5s and tRF-3s, from the extreme 5’ and 3’ ends of mature tRNAs respectively, and tRF-1s, from the 3’ trailer of precursor tRNA ((Lee et al. 2009); reviewed in (Keam and Hutvagner 2015; Kumar et al. 2016)). Moreover, recent studies indicate that 30-36 base long tRFs have essential roles in transgenerational gene regulation by metabolic stress: mice with high fat diet have higher levels of tRNA derived RNA fragments in their sperm and this set of tRFs serves as a signaling molecule in their offspring (Chen et al. 2016; Sharma et al. 2016). As a result, the offspring of high fat diet mice have a greater tendency to develop obesity and diabetes. These 30-36 base long fragments generated by cleavage in the anticodon loops are also called tiRNAs/tRNA halves and have been tied to stress response (reviewed in (Saikia and Hatzoglou 2015)). On the other hand, the roles of shorter tRFs have not been investigated in a detailed manner.

Different from the 30-36 base long tRNA halves, the shorter tRFs (18-22 bases long) are in the size range to enter Argonaute based on structural information (Elkayam et al. 2012; Schirle and MacRae 2012). Thus we first sought to bioinformatically test how tRFs associate with canonical miRNA machinery. A meta-analysis of available short RNA sequencing data suggested that although tRFs are produced in mouse embryonic stem cells that are *Dicer-/-,* their levels in Argonautes, the main effector proteins in the miRNA pathway, are as high as miRNAs (Kumar et al. 2014). Moreover, analysis of PAR-CLIP data (Hafner et al. 2010) indicated that tRF-3s and tRF-5s bind to Ago proteins in a manner similar to miRNAs and use their 5’ seed sequence of 7-8 bases to bring their target mRNAs into the Ago complexes (Kumar et al. 2014). Most intriguingly, CLASH data (Helwak et al. 2013) analysis identified more tRF-3-mRNA chimeras than miRNA-mRNA chimeras associated with Ago1 (Kumar et al. 2014), suggesting a potential function of tRF-3s in regulating gene expression using a mechanism similar to miRNAs.

To explore the role of tRF-3s, we devised ways to up-regulate endogenous tRF-3 levels. We discovered that overexpression of a particular tRNA (tRNA LeuAAG, tRNA CysGCA or tRNA LeuTAA) resulted in production of the specific tRF-3 (tRF-3001, tRF-3003 and tRF-3009, respectively) (nomenclature based on tRFdb (Kumar et al. 2015)). The tRF-3s specifically downregulate expression of luciferase reporters with a complementary sequence in the 3’ UTR. This repression is Dicer-independent but Argonaute dependent and also dependent on match to the seed sequence at the 5’ end of the tRF. tRF-3s associate with P-bodies based on the TNRC6 protein PAR-CLIP data analysis. Furthermore, RNA-seq of cells overexpressing the tRFs showed that mRNA targets of tRF-3s are significantly repressed. c-Myc oncogene is known to upregulate global tRNA expression (Lin et al. 2012). Consistent with the results with individual tRNA overexpression, c-Myc overexpression induced corresponding tRF-3s and repressed their targets globally. Thus, tRF-3s, generated independent of miRNA biogenesis proteins such as Dicer and Drosha, are able to enter into Argonaute complex to regulate gene expression via seed matches.

## Results

### Overexpression of tRNA results in production of the corresponding tRF-3

tRF-3s are 18-22 nucleotide small RNAs of two distinct sizes; tRF-3a (~18 nt) and tRF-3b (~22 nt) (Kumar et al. 2014) (Fig. 1A). We overexpressed tRNAs in HEK293T cells and assessed tRNA and tRF levels by QRT-PCR. The overexpression of tRNA LeuAAG, tRNA CysGCA and tRNA LeuTAA resulted in overproduction of the corresponding tRF-3s (Fig. 1B and 1C).

**Figure 1:**
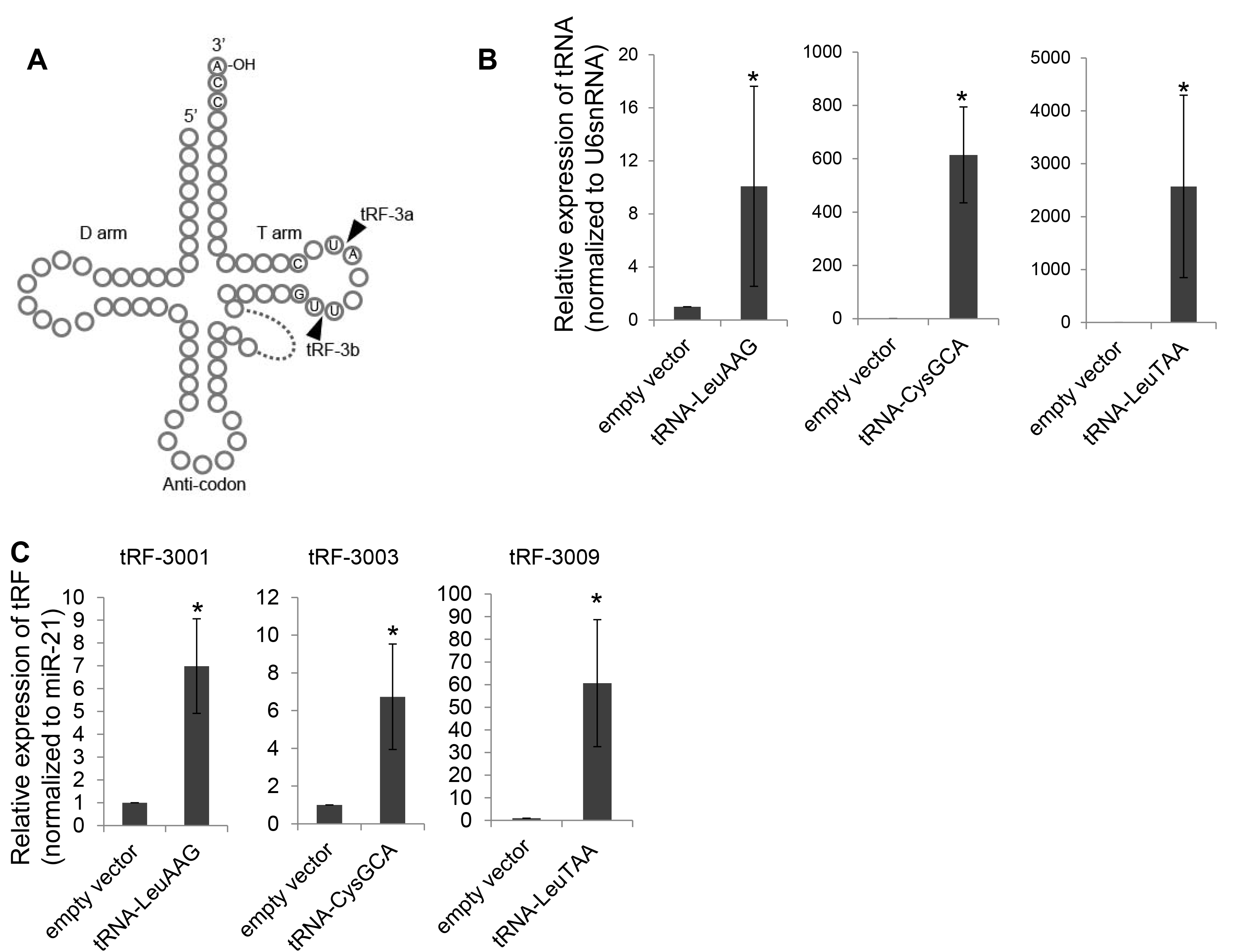
Production of endogenous tRF-3s by tRNA overexpression. (**A**) tRNA secondary structure depicting the tRF-3 cleavage sites. (**B**) Relative levels of indicated tRNAs upon tRNA overexpression. Mean and s.d. of three independent experiments. (**C**) Relative levels of indicated tRFs upon overexpression of the tRNAs indicated in (B) in the same order from left to right. Mean and s.d. of at least three independent experiments.

### tRF-3s silence gene expression through sequence complementarity

Luciferase reporters with perfect complementary sequences to the tRFs in the 3’ UTR were co-transfected with the tRNA. When tRF-3001 is expressed by overexpressing tRNA-LeuAAG and the luciferase 3’UTR has a perfect complementary sequence to tRF-3001, the luciferase activity is repressed to 40% (Fig. 2A). Luciferase reporters with complementarity to the cognate tRFs were down regulated similarly upon tRF-3003 and tRF-3009 overexpression (Fig. 2A).

**Figure 2:**
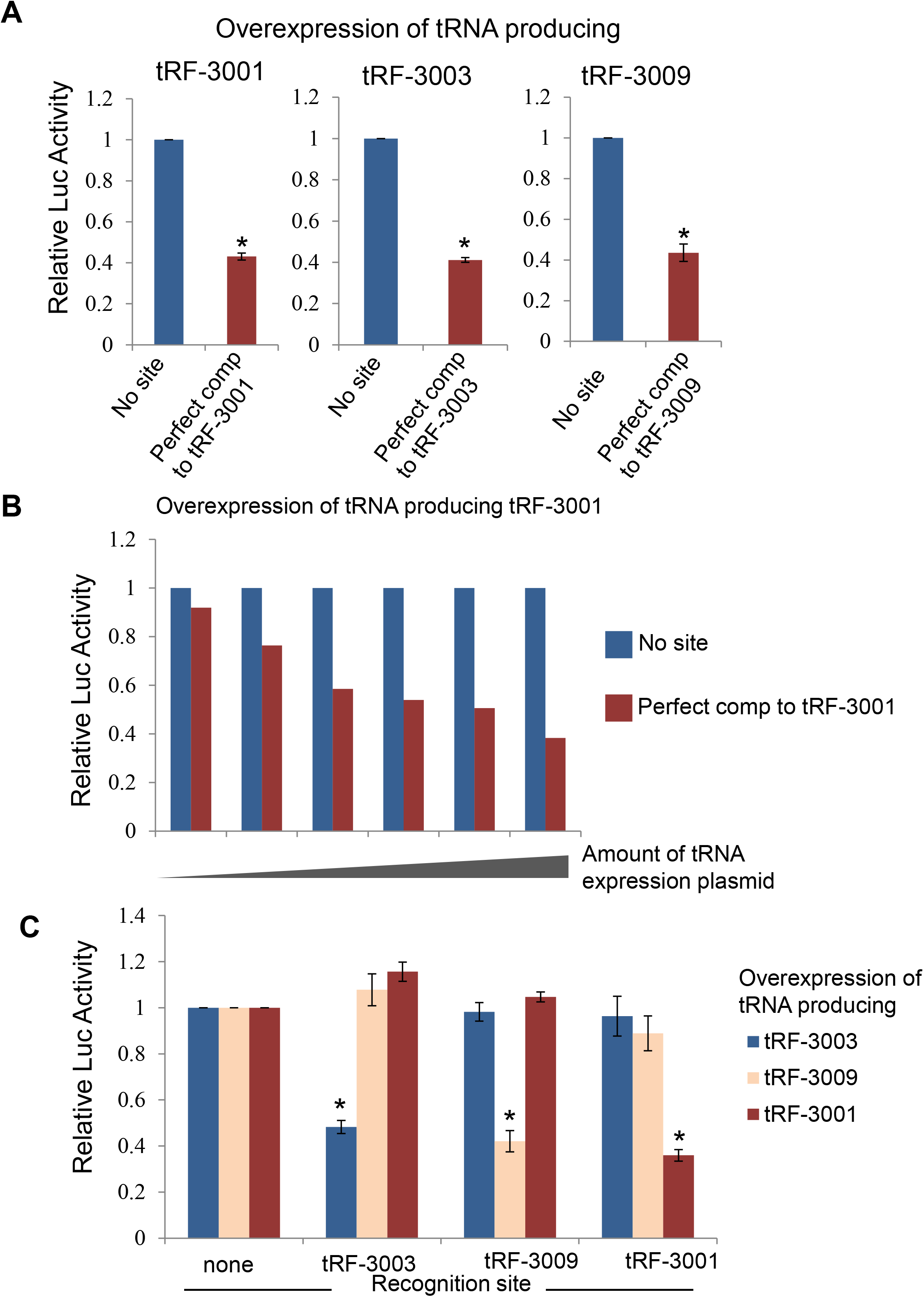
tRF-3s down regulate target expression through complementarity in 3’UTR of luciferase reporter. (**A**) Luciferase reporter assays using Renilla luciferase with a perfect complementary sequence to tRF-3 at the 3’UTR. Renilla luciferase levels were first normalized to Firefly luciferase levels from the same transfection and then normalized to no tRNA/tRF overexpression (empty vector) control. *: p-value <0.05. (**B**) Degree of repression correlates with tRNA overexpression amount. Amount of transfected tRNA plasmid was titrated to measure the change in the degree of repression. Luciferase reporter assays were analyzed as described in Fig. 2A. (C) Luciferase reporter assay showing the specific repression by each corresponding tRF-3. *: p-value <0.05

Titration of the amount of tRNA expressing plasmid showed that the downregulation of luciferase is directly correlated with the amount of the tRNA expressing plasmid (Fig. 2B and Supp. Fig. 1). We tested the specificity of the downregulation by transfecting tRNA expressing plasmid and luciferase reporter plasmid with complementarity to the other two tRF-3s and no repression was observed (Fig. 2C). For example tRF-3001 producing plasmid selectively repressed the luciferase reporter with complementarity to tRF-3001, but not those with complementarity to tRF-3003 or tRF-3009 and so on (Fig. 2C).

### tRF-3s have seed sequence properties similar to miRNAs

miRNAs interact with their target mRNAs by base pairing. In plants, perfect complementarity is essential for miRNA-mRNA targeting. In metazoans, with few exceptions, miRNA-mRNA base pairing occurs imperfectly allowing unmatched bulges between the two RNAs. The miRNA-mRNA base pairing rules are defined based on both bioinformatics and experimental results. The most important rule is that miRNAs recognize their mRNA targets using their 2-7 nt sequence at their 5’ ends, which is defined as the seed sequence. Second, an unpaired region at the middle of miRNA-mRNA pairing (bulge) is tolerated. Third, following the bulge there is often sequence complementarity between miRNA and mRNA ((Brennecke et al. 2005; Lewis et al. 2005; Grimson et al. 2007; Nielsen et al. 2007) and reviewed in (Filipowicz et al. 2008)). To test whether tRF-3s also follow these rules, we mutated the target site on the luciferase reporter three bases at a time.

As mentioned above, tRF-3s can have two distinct lengths: 18nt (tRF-3a) or 22nt (tRF-3b), and the short RNA sequence data analysis suggests that tRF-3003 and tRF-3009 are present in both “a” and “b” forms while tRF-3001 only has “a” form (Kumar et al. 2015). The luciferase reporters have perfect complementarity to tRF-3b so that both forms of tRFs could repress the target. Mutations that disrupted pairing of the target with the 5’ seed of each tRF-3a failed to repress the target (M1, M2, M3 in Fig. 3A and M3, M4 in Fig. 3B and M3 in Fig. 3C). The mutational analysis, however, showed that repression was mostly mediated via tRF-3a, because mutations at the extreme 3’ end of the target, that would disrupt pairing with the seed sequence of the longer tRF-3b, but not tRF-3a, continued to be repressed by the tRNA overexpression at similar level as perfect complementary sequence (M1, M2 in Fig. 3B and 3C).

**Figure 3:**
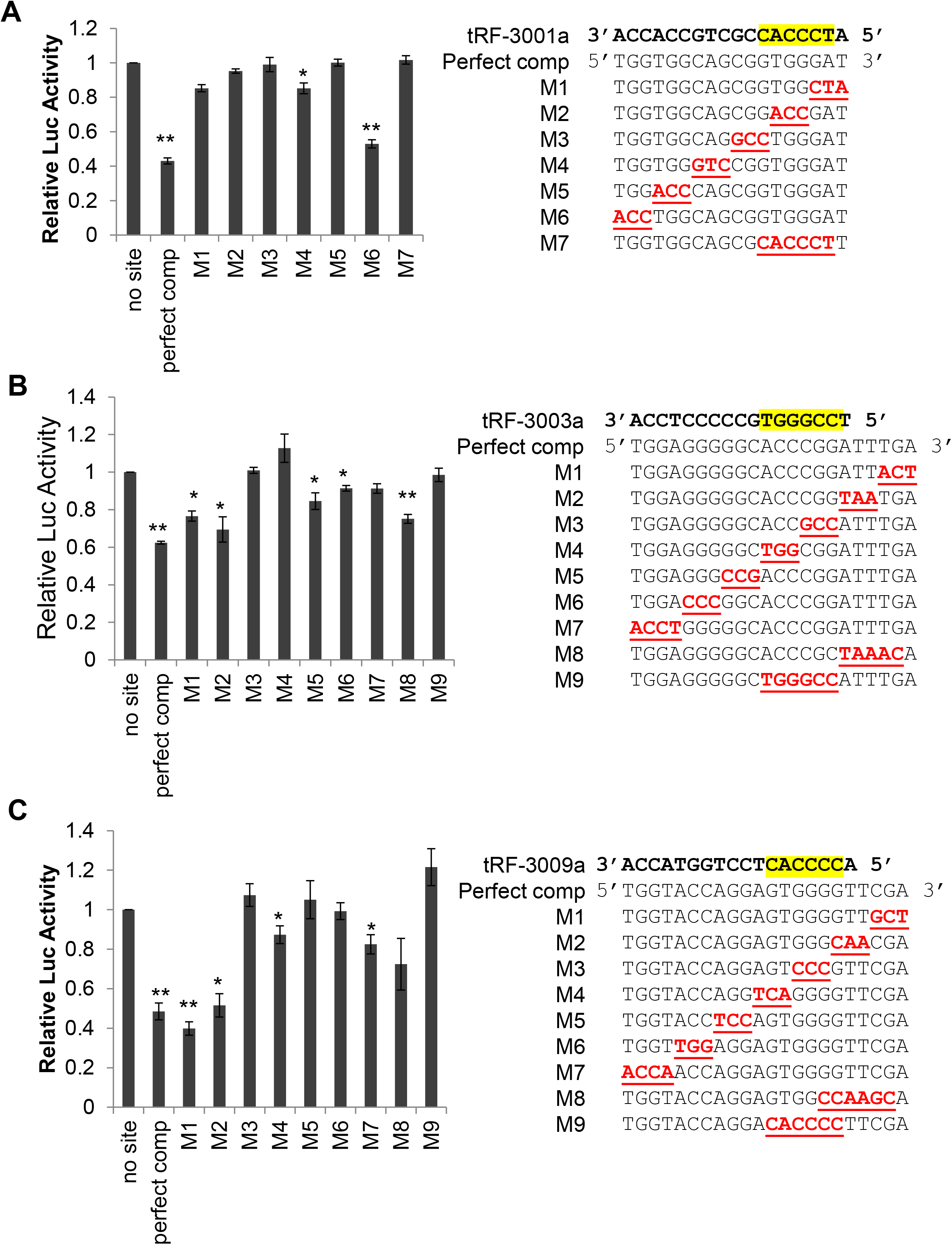
Seed sequence is required for target repression by tRF3s. Luciferase reporter assays with mutant target site at the luciferase reporter upon tRF-3001 **(A)**, tRF-3003 **(B)** and tRF-3009 **(C)** overexpression. Canonical seed region on each tRF-3 is highlighted in yellow. Mutated regions are underlined and colored red. P values are calculated by t-test comparing luciferase reporter with indicated target sequence to empty vector control (*: p-value < 0.05, ** : p-value < 0.005).

Perfect complementarity to the entire length of the tRF was not required for repression. For example, tRF-3003a can still repress a target that has mutations at the middle of the tRF-mRNA pair indicating tolerance for a bulge structure in the middle of the tRF-mRNA pairing (M5 in Fig. 3B). Mutations further away from canonical seed regions also allowed repression (M6 in Fig. 3A and 3B). In all cases, mutations outside of canonical seed regions (M5, M6 and M7 in Fig. 3B) showed lower degree of repression than with the perfect match target. A similar pattern is seen for tRF-3001 and tRF-3009: target with selected mutations 3’ to the seed sequence allowing some repression, but less than with perfect complementarity (M4, M6 in Fig. 3A and M7 in Fig. 3C). However these two tRFs appear to require additional complementarity outside the seed sequence for repression (e.g. M5 in Fig. 3A and M5, M6 in Fig. 3C). Thus the rules of repression are similar to microRNAs, in that the target should be complementary to the 5’ seed sequence of the tRF and though perfect complementarity to the entire length of the tRF is not essential for repression, additional complementarity downstream from the seed sequence facilitates repression.

### Gene silencing through tRF-3 is independent of Dicer but dependent on Argonaute proteins

One of the major enzymes in miRNA pathway is Dicer, which cuts out the loop in the stem-loop hairpin structure of the precursor miRNA (pre-miRNA) to generate miRNA (Bernstein et al. 2001). Analysis of small RNA sequencing data from Dicer knock-out cell lines (Bogerd et al. 2014) showed that although Dicer is required for microRNA biogenesis, it is not required for the biogenesis of tRF-3001, tRF-3003 or tRF-3009 (Supp Fig 2). Consistent with this, the luciferase repression by tRF-3 is still observed in the *Dicer* knock-out cells, and is even more pronounced than in wild type cells (Fig. 4A), most probably because more Argonaute protein become available due to the failure to produce cellular miRNA.

**Figure 4:**
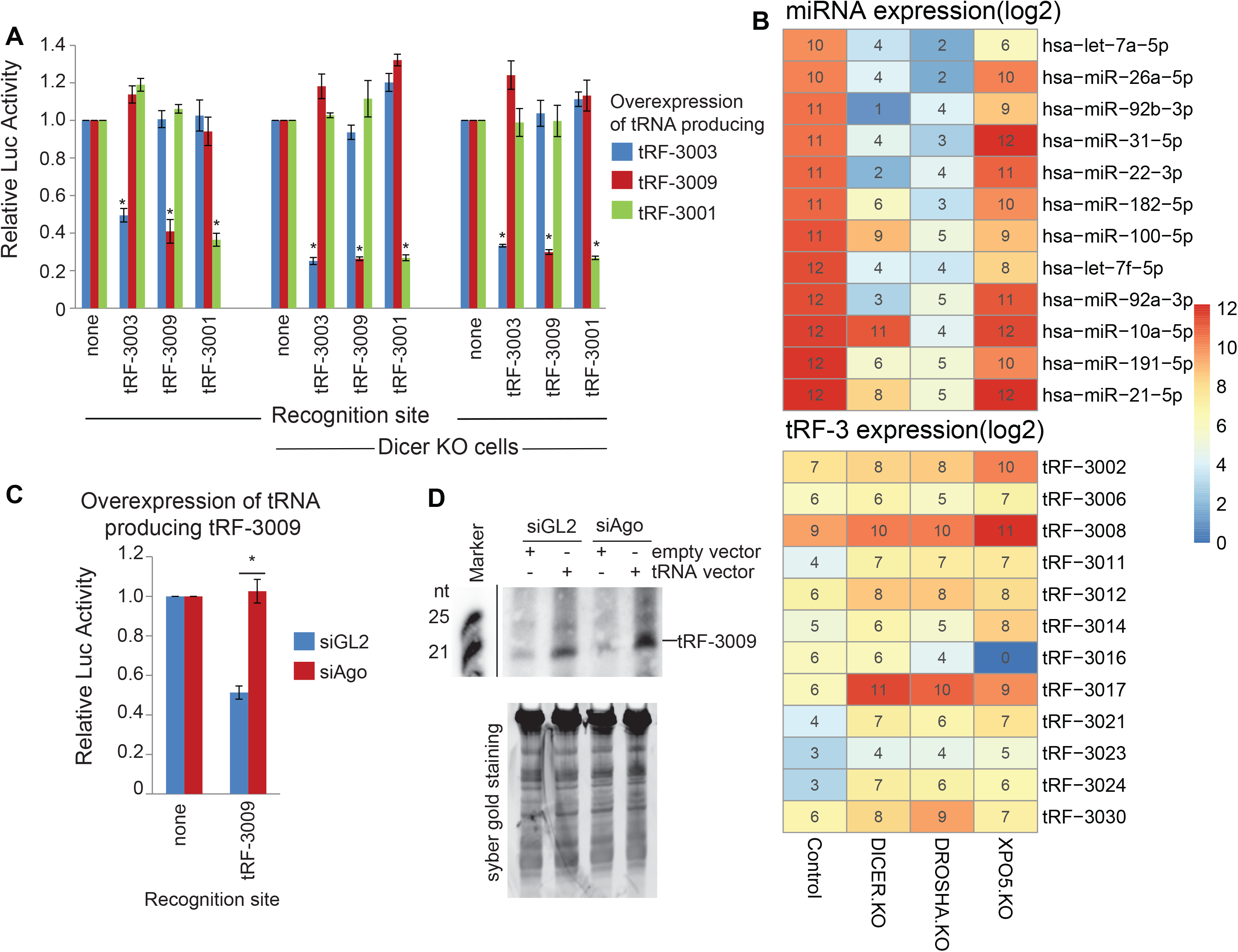
Target repression by tRF-3s is independent of Dicer but dependent on Argounates. **(A)** Luciferase reporter assays in wild type and Dicer knock out HEK293T cells, NoDice 4-25 (middle) and NoDice 2-20 (right) (Bogerd et al. 2014). **(B)** miRNA and tRF-3 read counts in small RNA sequencing data from WT, Drosha, Dicer and Exportin-5 knockout HCT116 cells (Kim et al. 2016). **(C)** Luciferase reporter assays after tRNA overexpression to produce tRF-3009 ± knockdown of Argonaute proteins. **(D)** Northern blot showing tRF-3009 levels in tRNA overexpressing cells after Ago knock down. Lower panel shows equal loading of lanes.

Our previous report showing that tRF abundance was unchanged in the absence of Dicer or DGCR8 was derived from embryonic stem cells (Kumar et al. 2014). Since then small RNA sequencing data has become available from *Dicer-/-, Drosha-/-* and *Exportin 5-/-* somatic cells (HCT116) generated using CRISPR-Cas9 technology (Kim et al. 2016). Analysis of this data showed that although Dicer and Drosha are important for microRNA levels, they are dispensable for tRF-3 generation (Fig. 4B). Exportin 5 depletion did not affect tRF-3 levels (Fig. 4B), similar to previous observations that Exportin 5 is dispensable for miRNA biogenesis (Kim et al. 2016).

To test whether Argonaute proteins (Ago) are essential for the gene repression function of tRF-3s, we repeated the luciferase reporter assays with Ago-1, -2 and -3 levels down regulated by siRNA transfection. Ago-4 levels also decreased upon knock down of Ago1, 2 and 3 (Supp Fig. 3A). As seen in Fig. 4C, the repression of the luciferase reporter by tRF-3009 was diminished when the Argonaute proteins were down regulated. Repression by tRF-3001 or tRF-3003 was also attenuated in response to lower Argonaute protein levels (Supp. Fig. 3B & 3C). By northern blot analysis, tRF-3009 is still generated as efficiently as in WT cells after Ago knockdown (Fig. 4D). Therefore, Ago is necessary for gene repression by tRF-3s but not for their biogenesis.

### tRF-3s interact with P-body proteins

When miRNAs interact with their target mRNAs in Ago proteins the pair of RNAs are often directed to P bodies, which leads to degradation of the target mRNA (Liu et al. 2005a; Liu et al. 2005b). To test whether tRF-3s also direct their targets to P bodies, we investigated the association of tRF-3s and P-bodies by analyzing PAR-CLIP data from HEK293 cells (Hafner et al. 2010). As seen in Fig. 5A, TNRC6A/B/C, components of the P bodies, associate with some tRF-3s, but not all. The most abundant association was observed for TNRC6A, in which case a few tRF-3s are present at levels comparable to miRNA levels (>1000 RPM), although in general tRF-3s associate with TNRC6 proteins at a lower level compared to microRNAs. Moreover, the locations of the T to C mutations in the PAR-CLIP data indicate that tRF-3s directly interact with TNRC6 proteins at position 12 and 14, while sparing bases 1-6 that overlap with the seed (Fig. 5B). This interaction is reminiscent of miRNA-TNRC6 interaction (Hafner et al. 2010), at position 11 and 13, with the bases 1-6 in the seed sequence being spared from the cross-link.

**Figure 5:**
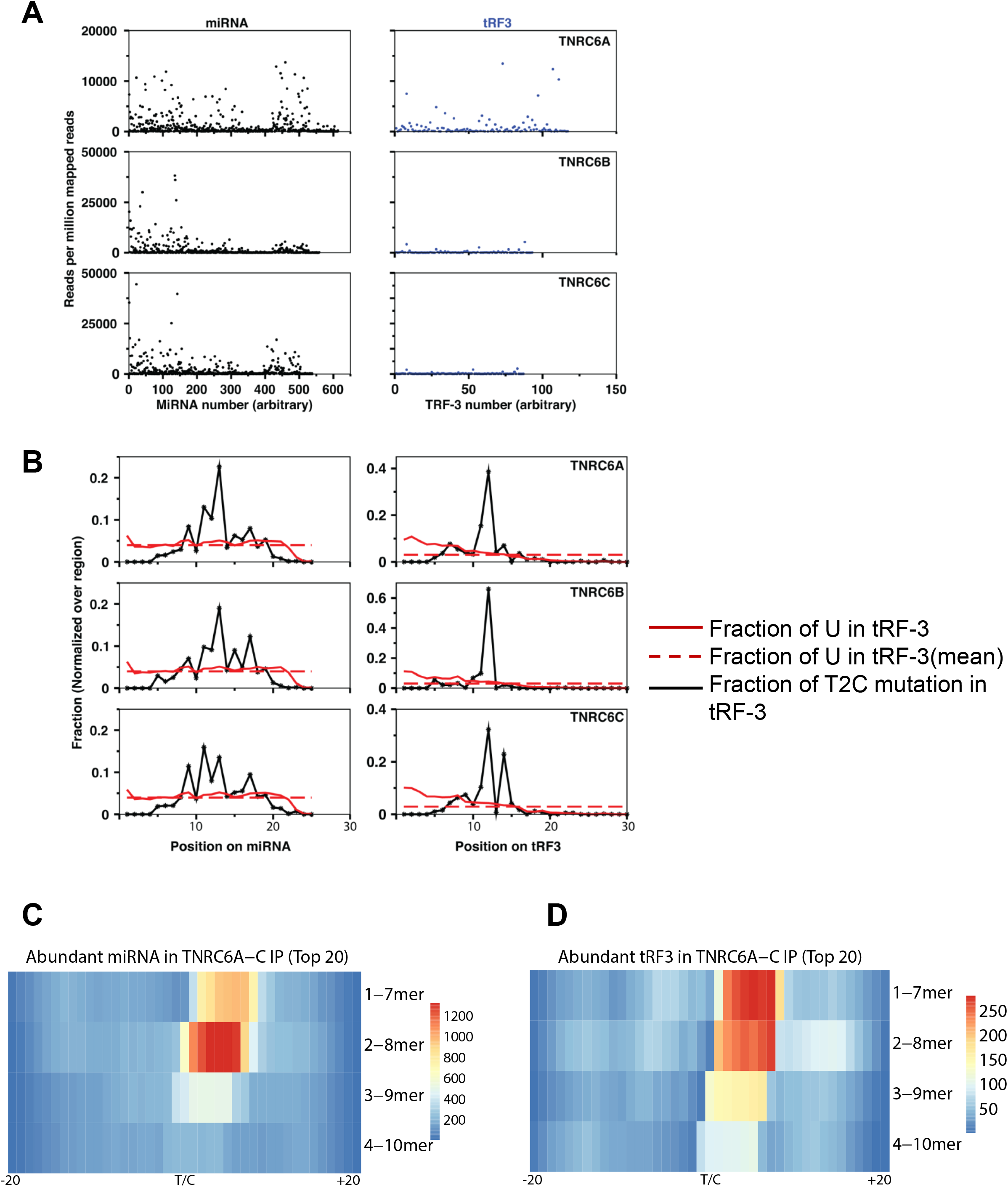
tRF-3s associate with TNRC6 proteins. **(A)** Read counts for miRNAs or tRF-3s in TNRC6A, TNRC6B and TNRC6C PAR-CLIP data from (Hafner et al. 2010). Each microRNA or tRF is given an arbitrary identifying number along the X-axis. The expression level (expressed as reads per million mapped reads) of each microRNA or tRF is shown on the Y-axis. **(B)** Normalized positional T to C mutation frequencies for miRNA or tRF-3 reads found in the TNRC6A, TNRC6B and TNRC6C PAR-CLIP data. **(C-D)** Complementarity in the target RNA CCRs present in the TNRC6A PAR-CLIP to the 1-7, 2-8 3-9 and 4-10-mer sequences from 5’ end of 20 most abundant microRNAs or tRF-3s seen in (B). The CCRs are centered on the site of the U/C mutation and the number of targets with complementarity indicated at the corresponding base in the sequence of the CCR. For example, the maximum number of matches is seen with the 1-7 mer of the tRF and begins with the base immediately downstream from the cross-link site in the target RNA.

The absence of any cross-link of the TNRC6 with the seed sequence of the tRF-3 or microRNA is likely because the seed is paired with the target RNA even in the TNRC6 complexes. Although the targets are degraded quickly in P bodies, we wondered whether a small fraction of microRNA or tRF-3 targets may be detected in the TNRC6 PAR-CLIP data. We therefore searched in the PAR-CLIP data for target RNA reads with complementarity either to seeds of microRNAs, or specifically to tRF-3s but not microRNAs. Targets that were detected were aligned with the T to C mutation marking the cross-link at the center along with 20 bases up and downstream from the cross-link to produce cross-link-centered-RNAs (CCRs). Target RNAs complementary to microRNAs can be detected in the CCRs associated with TNRC6 (Fig. 5C). Most interesting, target RNAs with complementarity to tRF-3 seeds are also associated with the TNRC6 (Fig. 5D). In the case of both the microRNAs and the tRF-3s, the most frequent complementarity to the seed is seen just downstream from the cross-link center, similar to what is seen with the microRNA:target or tRF-3:target pairs in Ago1-4 PAR-CLIP data (Hafner et al. 2010), suggesting that similar pairs of RNAs are transferred to P bodies.

### RNA-seq analysis reveals that tRF-3s repress endogenous mRNA targets globally

miRNA-mRNA loaded RISC complex mediates the degradation of mRNA targets with the help of P bodies. Since tRF-3s paired with their target RNAs are also found in P bodies, we decided to analyze gene expression changes by RNA-sequencing upon tRNA/tRF overproduction. As seen in Figure 6A, upon overexpression of tRNA CysGCA which produces tRF-3003, target mRNAs whose 3’UTRs bear complementarity to 8mer, 7mer-A1, 7mer-M8 and 6mer seed sequences are repressed relative to those that do not have such complementarity in their 3’UTR. Similar to miRNAs, targets complementary to the full 8mer are repressed the best, followed by those complementary to 7mer-A1, 7mer-M8 and 6mer seed sequences. tRF-3009 also repressed its targets with similar gradation of efficiency for matches to each seed-mer (Fig. 6B). These experiments were repeated again and cumulative distribution function plots of the second replicate can be found in Supp. Fig. 4.

**Figure 6:**
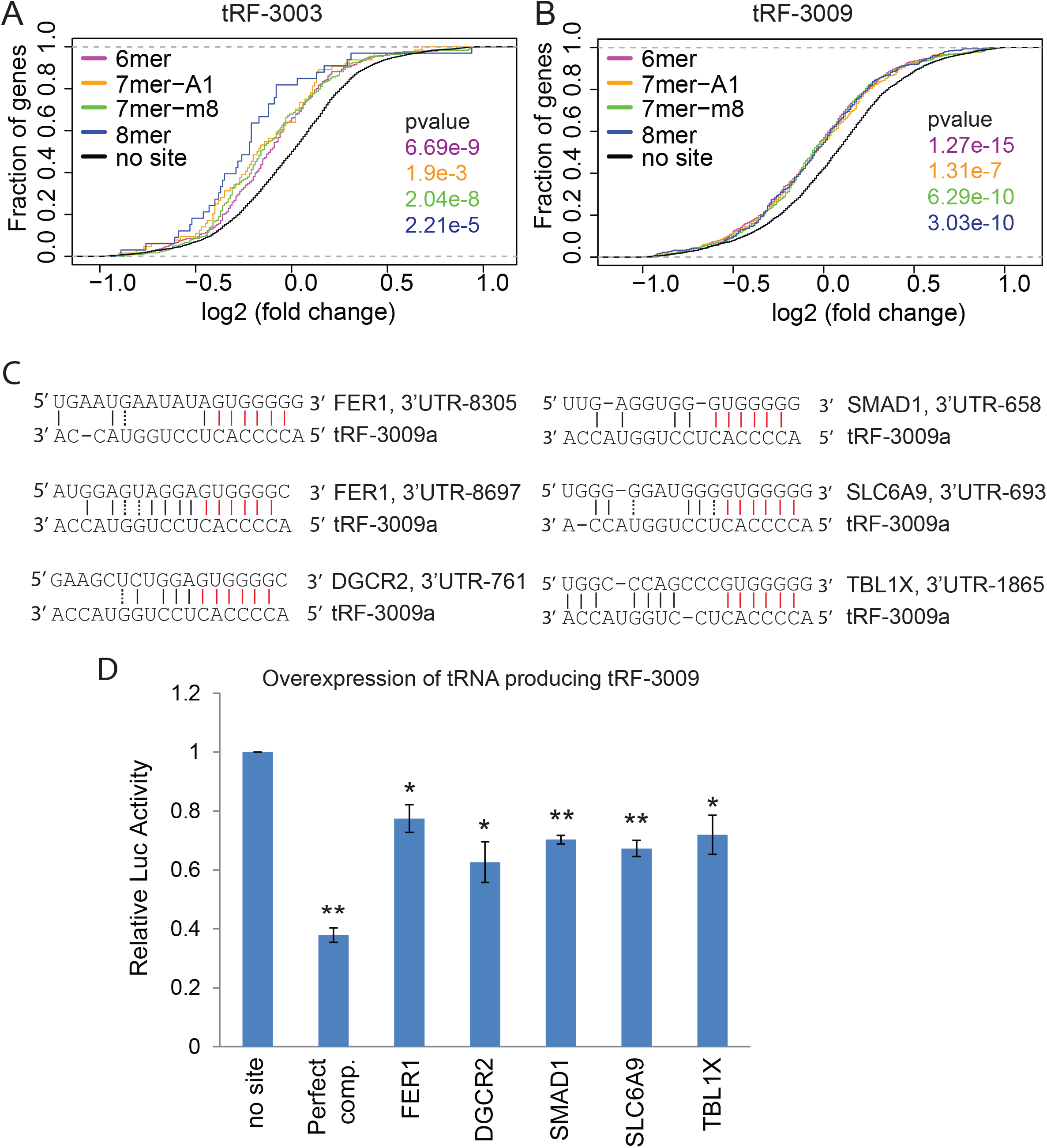
Global target repression by tRNA/tRF overexpression is mediated through seed complementarity. Cumulative distribution function (CDF) plots showing the repression of tRF-3 targets upon overexpression of tRNAs producing tRF-3003 **(A)** or tRF-3009 **(B)**. Targets are classified in a similar way to miRNA targets. **(C)** Seed sequence complementarity in selected tRF-3009 targets that are identified in RNA-seq upon tRF-3009 expression. The number after 3’UTR indicates the start position of the mRNA sequence match with 1 being the base immediately downstream from the stop codon. Red line: perfect base-pairing in seed; black line: perfect base-pairing outside seed; dashed line: wobble base-pairing. FER1 gene has two predicted complementary sites on its 3’ UTR. **(D)** Dual luciferase reporter assay on reporters containing the 3’UTR of indicated genes after tRNA-LeuTAA transfection, leading to tRF-3009 overexpression. Perfect complementary sequence to the tRF serves as a positive control and all results are normalized to the “no site” reporter without any match to the tRF (*: p-value < 0.05, **: p-value, 0.005,).

To validate the putative tRF-3 targets, we picked the top 5 repressed targets after tRF-3009 overexpression and performed luciferase reporter experiments with corresponding 3’UTR sequences from these targets. The 3’UTR sequence from these repressed endogenous targets have base complementarity with tRF-3009 seed region (Fig. 6C). Luciferase reporters containing the 3’UTR of these endogenous genes are all repressed upon overexpression of tRNA-LeuTAA producing tRF-3009 (Fig. 6D). This result independently validated the repression observed in the RNA-seq experiment.

### Myc-overexpression induces tRF-3s and represses their targets through seed complementarity

There is an increased occupancy of c-Myc at the promoters of actively transcribing genes, which results in global induction of nearly all genes in tumor cells that overexpress Myc (Lin et al. 2012). c-Myc also occupies the promoters of rRNA and tRNA genes, and overexpression of Myc is known to induce tRNA genes (Gomez-Roman et al. 2003; Arabi et al. 2005; Grandori et al. 2005; Lin et al. 2012). Since overexpression of tRNA induced the levels of tRF-3s (Fig. 1C), we first checked the levels of four tRFs upon overexpression of Myc for 24 hrs in P493-6 B-cell lymphoma cells (Lin et al. 2012; Loven et al. 2012). The levels of tRF-3007, tRF-3006, tRF-3002 and tRF3003 are elevated upon Myc induction. (Fig. 7A-C respectively and Supp. Fig. 5). Most importantly, the predicted targets of the tRF-3s were repressed compared to non-targets when Myc is induced (Fig. 7D-F). Stratification of substrates based on complementarity to different types of seed sequences; again suggest that 8-mer seed match targets are mostly repressed better than other types of seed match targets (Fig.7D and F). The 7mer seed match targets are repressed nearly as well as the 8mer seed matches. These results suggest that under conditions where tRNAs are naturally induced, the resulting co-induction of tRFs modulates the gene expression pattern.

**Figure 7:**
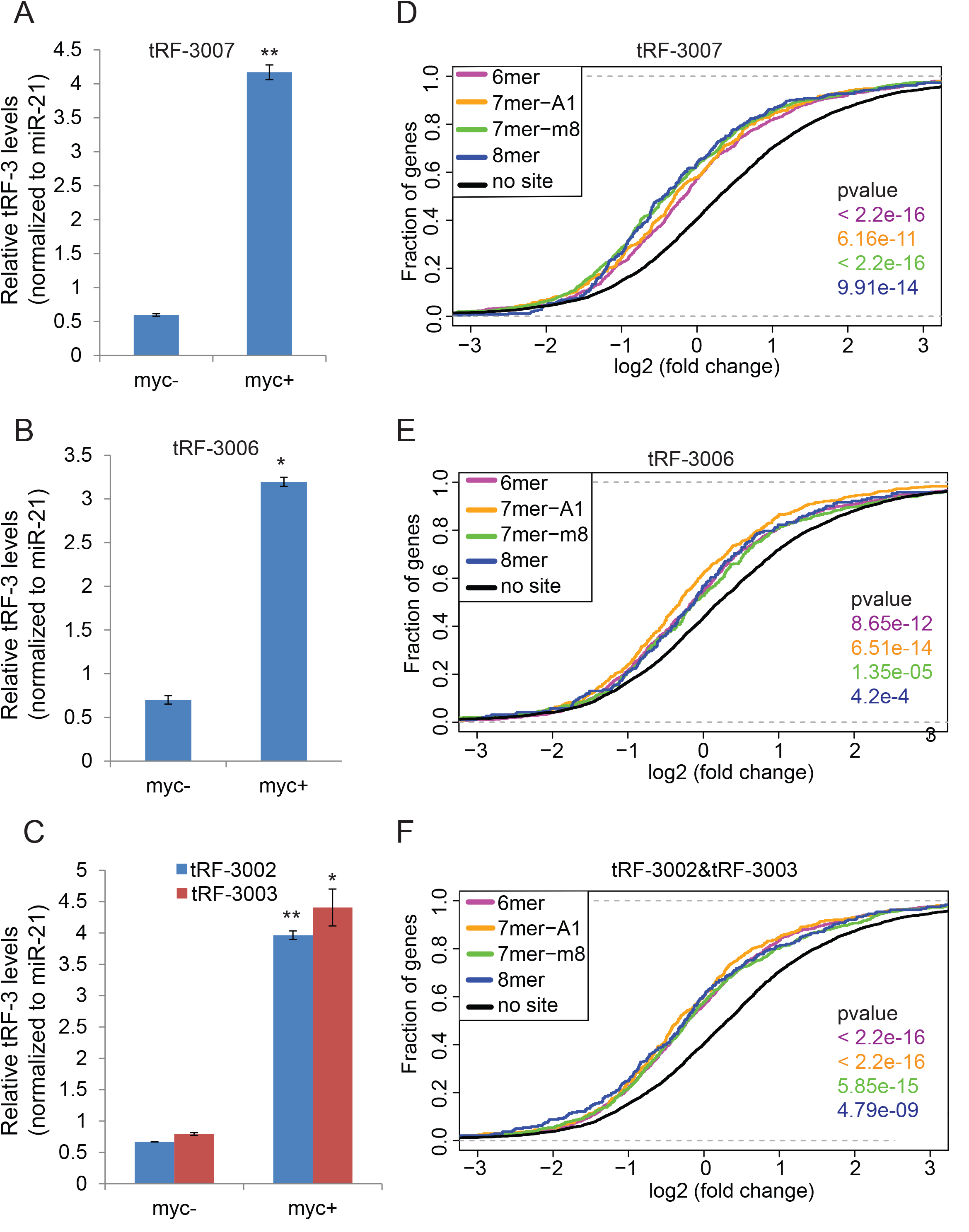
tRF-3 targets are repressed through seed complementarity upon Myc overexpression. **(A-C)** Up-regulation of endogenous tRF-3s upon Myc overexpression in P493-6 cells. Relative levels of tRF-3007, tRF-3006, tRF-3002 and tRF-3003 are quantified by RT-QPCR after size selection (myc- : 0 hr; myc+: 24 hr after Myc induction). Mean ±s.d. of 3 technical replicates. (Another independent experiment is shown in Supp. Fig. 5). *: p-value < 0.005, **: p-value < 0.0005). **(D-F)** Cumulative distribution function (CDF) plots showing global repression of targets by tRF-3007, tRF-3006 and tRF-3002/tRF-3003 upon Myc overexpression. Targets of tRF-3s are predicted based on seed complementarity. tRF-3002 and tRF-3003 have the same seed sequence.

## Discussion

Recent advances in sequencing technology have led to the discovery of different small RNA fragments including tRNA derived RNA fragments. Here we show that tRF-3s, which are generated from the 3’ part of the mature tRNA down-regulate target gene expression in a sequence dependent manner similar to miRNAs. Most importantly, our RNA-seq analysis showed that tRF-3s can regulate gene expression in a global manner. To the best of our knowledge, this is the first report showing that tRF-3s that are generated by overexpression of their parental tRNAs can down-regulate gene expression globally. miRNAs repress their targets most effectively if there is an 8-mer seed match in the target (Lewis et al. 2005; Grimson et al. 2007; Hafner et al. 2010). These observations were also supported by the structure of human Ago2 complexed with a miRNA and its target (Schirle et al. 2014). Both our mutational analysis on the luciferase reporter and RNA-seq analysis after tRNA overexpression show that similar seed pairing rules are also important for tRF-3s to recognize and repress their targets.

Oncogene Myc was shown to upregulate PolIII transcripts including tRNAs (Gomez-Roman et al. 2003; Arabi et al. 2005; Grandori et al. 2005; Lin et al. 2012). Here we showed that tRNA upregulation after Myc overexpression results in upregulation of tRF-3s. Strikingly, the targets of several tRF-3s are repressed upon Myc overexpression.

The analysis of publicly available short RNA sequences from *Dicer* knock out mouse embryonic stem cells showed that Dicer is not required for generation of tRF-3s (Kumar et al. 2014; Kumar et al. 2015). In contrast, there are papers reporting that Dicer might be necessary for tRF generation (Cole et al. 2009; Maute et al. 2013). We find, however, that in multiple cell lines tRF-3 generation is Drosha- and Dicer-independent and yet Argonaute proteins are essential for their function (Fig. 4). Remarkably, tRF-3s represses their targets better in *Dicer* knock out cells suggesting that Argonaute proteins are more accessible for tRF-3 loading in the absence of miRNAs (Fig. 4A). This result raises the possibility that tRFs become the main regulators of gene expression in the absence of miRNAs. Downregulation of Dicer and/or Drosha have been correlated with a worse outcome in lung, breast, skin, endometrial and ovarian cancer (reviewed in (Foulkes et al. 2014)). Our results suggest that tRF-3s acquire more potency when there are lower levels of Dicer or Drosha so that we should investigate whether tRFs are important for the poorer outcome in these tumors.

On the other hand, Dicer has an important role in loading microRNAs into Argonaute complexes (Chendrimada et al. 2005; Gregory et al. 2005). The fact that tRF-3s can repress genes by a mechanism dependent on Argonaute proteins in Dicer knock out cells suggest that Dicer is not essential for loading of all short RNAs into Argonaute. It is possible that the Dicer-independent loading of tRF-3s into Argonaute is less efficient than the Dicer-dependent loading of microRNAs. However, our results opens the possibility that other short RNAs, if present at high concentration, may load on to Argonaute complexes and repress gene expression. It has been shown that Hsp90 proteins help the stability of unloaded Argonaute proteins and loading of miRNA to Argonaute in an ATP-dependent way (reviewed in (Meister 2013)), and thus Hsp90 may be involved in the loading of tRFs on Argonaute proteins by this Dicer-independent pathway. Recently agotrons, short introns interacting with Argonaute proteins, were shown to be loaded into Argonaute independently from Dicer, suggesting that such loading is not unique to tRFs (Hansen et al. 2016).

The lack of any requirement for Dicer or Drosha for tRF-3 production opens up the question of how tRF-3s are generated. Identification of the specific enzyme/enzymes that are important for tRF generation will let us better understand tRF biology and function.

Similar to miRNAs, some tRF-3s directly interact with GW proteins in P bodies and decrease levels of target mRNAs. But clearly, many tRF-3s do not do so, and in those cases changes in target protein levels will have to be sought to determine whether they too repress gene expression. As with microRNAs, the reason for the differential interaction of specific tRF-3s (and their targets) with P bodies need to be elucidated.

Finally, it is very exciting that in Myc transformed cells we see an induction of tRF-3s and a corresponding repression of the targets of tRF-3s. This shows that even natural induction of tRNAs can regulate gene expression through the induction of tRFs. The results here will be germane to the action of many other oncogenes known to induce tRNAs. Examples include Ras, Raf, EGF receptor and oncogenes like E6 and E7 from human papilloma virus (that inhibit p53 and Rb, both known to repress tRNA transcription) (Grewal 2015).The results here will also be relevant for other growth stimulatory pathways associated with tRNA induction, e.g serum stimulation.

In summary, tRF-3s, abundant short RNA fragments derived from tRNAs can regulate gene expression by mechanisms similar to microRNAs. These results expand the repertoire of endogenous short RNAs that are in use for regulation of gene expression.

## Material and Methods

### Cloning

Synthetic oligonucleotides containing perfect complementarity to tRF-3s were cloned at the 3’UTR of Renilla luciferase gene in psi-CHECK2 (Promega) reporter plasmid by infusion cloning between PmeI and XhoI sites for luciferase assays. 3’UTRs of endogenous genes were similarly cloned into psi-CHECK2.

### RT-qPCR analysis of tRNAs/tRF-3s

4 μg of tRNA overexpression plasmids were transfected into HEK293T cells in 6 cm plates using Lipofectamine 2000 (Thermofisher). The cells were collected after 2 days of transfection and total RNA was purified for further analysis.

For detection of tRNAs by RT-qPCR, total RNAs were purified with TRIzol (Ambion) extraction. 1μg of total RNA was subjected to DNase treatment using RQ1 RNase free DNase from Promega. cDNAs were generated using Super script III reverse transcriptase (Thermo Fisher Scientific) using random hexamers as primers according to manufacturer’s instructions. Quantitative PCR were performed using Sybr green mixes and specific primers against tRNA. Levels were normalized to U6snRNA gene.

For detection of tRFs, 100 μg of total RNA was subjected to small RNA enrichment using miRVANA miRNA purification kit (Ambion). Small RNA enriched RNA pool was loaded into 15% 7M Urea-Polyacrylamide page and RNAs in the 15-35 nt size range were purified. The size selected RNA was eluted from gel slices overnight in 0.3 M NaCl/TE buffer at room temperature and precipitated with 2 volumes of isopropanol + 10 μg glycogen at −20°C overnight. RNA was precipitated by centrifugation at 4°C for 20 min at 13,000 rpm and washed once with 70% EtOH. cDNAs were generated using NCode miRNA First-Strand cDNA Synthesis and qRT-PCR Kits with 50-100ng 15-35 nt long RNA as starting material. The levels of tRFs are normalized to the levels of miR-21.

### Northern blotting

Cells were transfected and collected as described above.

40 μg of TRIzol extracted total RNA was loaded into 15% 7 M Urea-polyacrylamide gels. The RNA was transferred onto Hybond-N+ positively charged nylon membranes (GE healthcare, USA Cat No:RPN203B) using a Bio-Rad transblot apparatus at 200 mA (9 −10 V) for 3 hours (Bio-Rad, USA). The transferred RNA was cross-linked to the membrane by UV-irradiation for 1 min (Stratalinker, Stratagene) with 254-nm bulbs with autocrosslink option). The membrane was dried between two dry 3M papers by baking at 50°C for 30 min. The membrane was stored at 4°C between filter papers until use. The rest of the protocol was performed as described in (Huang et al. 2014). Briefly, membranes were pre-hybridized for at least 30 minutes at 40°C in pre-hybridization buffer (7% SDS, 200 mM Na2HPO4 (pH 7.0)) containing 5 μg/ml denatured salmon sperm DNA. Hybridization was performed in Expresshyb solution, Clontech Cat No: 636831 containing 50 pmol/ml labeled probes (Anti-3009: BIO-TGGTACCAGGAGTGGGGT) at 40°C overnight (typically 16 hrs). The membrane was washed thrice with 1X SSC, 0.1 % SDS at RT for 5 min. ECL Lightning was performed using Chemiluminescent Nucleic Acid Detection Module Kit from Thermo Scientific, USA following the instructions in their manual.

### Dual Luciferase assay

Luciferase assays were performed in 24-well plates. 480 ng of tRNA overexpression plasmid or empty vector pcDNA3 and 2 ng of psi-CHECK2 reporter vector with no sites or perfect complementary sites to tRF-3 in the 3’UTR of Renilla luciferase gene were transfected in to HEK293T cells using Lipofectamine 2000 (Life Technologies) as a transfection reagent. The cells were lysed in 100 ul 1X passive lysis buffer from Dual-Luciferase^®^ Reporter Assay System (Promega) after 2 days of transfection. The luciferase signals were measured using 20 ul of the lysate following the instructions provided by Promega. The Renilla luciferase levels were normalized to firefly luciferase levels and the results were always plotted with tRF overexpression over non-overexpression (see y axis on plots: tRF+/tRF-).

Luciferase reporter assays were done in the same way with reporters containing 3’UTRs of endogenous genes.

### siRNA knock down experiments

To knock down Argonaute proteins, 20 nM of siGL2 or 20 nM total of siRNAs against Ago1, 2 and 3 were transfected into HEK293T cells using RNAimax transfection reagent from Life Technologies, and this was repeated after 24hrs. Luciferase reporters and tRNA overexpression plasmids were transfected 24 hrs after second siRNA transfection. Lysates were collected 48 hr after second siRNA transfection for western blot analysis and dual luciferase assay.

### Western blotting

Argonaute protein levels were detected by Western blotting. Briefly, the membrane was incubated in 5% milk in TBS-T blocking solution for 1 hr at room temperature and in rabbit monoclonal primary (Ago1: CST#5053, Ago2: CST#2897) in 5% BSA, TBS-T with 1:1000 dilution for overnight at 4°C. The membrane was washed 3 times in 1X TBS-T and incubated with 1:5000 diluted HRP goat-anti-rabbit secondary in 5% BSA, TBST at room temperature. Immobilon Western Chemiluminescent HRP substrate from Millipore was used for developing the signal.

Myc induction in P493-6 cells was performed as described in (Loven et al. 2012). Protein levels were detected as described in (Blum et al. 2015).

### RNA-seq library preparation

HEK293T cells were transfected with empty vector or tRNA expression plasmids 4 times in two days intervals. Total RNA was purified with Qiagen RNeasy kit. tRNA and tRF levels were quantified as described above.

1 ug of total RNA from pcDNA3, tRNA-CysGCA (tRF-3001), tRNA LeuAAG (tRF-3003) and tRNA LeuTAA (tRF-3009) overexpressing cells were used for library preparation. Libraries were prepared using NEBNext Ultra directional RNA library prep kit for Illumina. The libraries were indexed using NEBNext Multiplex oligos for Illumina. Quality controls and sequencing was done in Genomic Services Lab at Hudson Alpha.

### Analysis of the small RNA data isolated from Dicer knock out and Dicer WT cells

We analyzed two small RNA datasets isolated from Dicer knock out and wild type cell lines generated by two independent laboratories (Bogerd et al. 2014; Kim et al. 2016). Firstly, adaptor sequence was removed using the `Cutadapt’ program (Martin 2011) and sequencing reads that were >=14 bases long were retained. To identify the total mapped reads the small RNA reads were mapped on whole genome (hg38 genome build) by using short read aligner Novoalign (http://www.novocraft.com/products/novoalign/). Next, the identical reads were collapsed and mapped on to the in house built small RNA database (mature miRNA and tRNA as detailed in (Kumar et al. 2014; Kumar et al. 2015)) using BLASTn (Altschul et al. 1990). The building of in house blast database for blast searches is explained in detail in our earlier publications (Kumar et al. 2014; Kumar et al. 2015). The total number of mapped reads was used for normalizing the expression of miRNA and tRFs. In general we considered only those alignments where the query sequence (small RNA) was mapped to the database sequence (tRNA or miRNA) along 100% of its length. The blast output file was parsed to get information on the mapped position of small RNA on tRNA or miRNA. We extracted all map positions where the small RNA aligned from its first base to the last base with the tRNA sequence allowing either one or no mismatch. Since ‘CCA’ is added at the 3’ end of tRNA by tRNA nucleotidyltransferase during maturation of tRNA (Xiong and Steitz 2006), we allowed a special exception for the small RNA mapping to the 3’ends of tRNAs allowing a terminal mismatch of <=3 bases. To remove any false positives, the small RNAs that mapped on to the ‘tRNAdb’ were again searched against the whole genome using blast search excluding the tRNA loci.

### Analysis of PAR CLIP data

We also investigated tRF and miRNA expression in human small RNA PARCLIP data of TNRC6A, B & C by analyzing the human TNRC6A (GEO ID = GSM545218), B (GEO ID = GSM545219), and C (GEO ID = GSM545220) PAR-CLIP data isolated from HEK293 cell lines(Hafner et al. 2010). Data from all three small RNA libraries were examined for miRNA and tRF expression as well as for the T to C mutation position and its frequency compared to wild type small RNAs (miRNAs and tRFs). Sequence reads that either mapped perfectly on miRNA or tRFs or mapped with one base mismatch were considered for T to C mutation analysis. Mismatched base and its position relative to the 5’ end of small RNA were collected for final analysis.

The 17,319 crosslink-centered regions (CCRs) identified by Hafner et al (Hafner et al. 2010) present in the PAR-CLIP data was used to study the complementary sequence of miRNA and tRF-3 seed sequence along the length of CCR. All the possible 7-mer sequence was generated along the length of miRNA and tRF. The 7-mer sequences were reverse complemented and mapped and the match was scored along the length of CCR. CCRs are 41 nt long sequences centered at the T (protein binding site) that showed the highest T to C frequency. Hafner et. al demonstrated that the reverse complement of known miRNA seeds is enriched in CCRs directly following this central cross-linked T. In our analysis four 7-mer (1-7, 2-8, 3-9 & 4-10) reverse complementary sequence of top 20 abundant miRNA and tRF-3 sequences identified in TNRC6A-C (a member of GW-bodies or P-bodies) were used for finding sequences along the length of CCR. The counting and scanning of the CCRs was done from the 5’ end to 3’ end of CCRs. Whenever there was a match, the score (count) was assigned to all the seven bases of CCR.

### Analysis RNASeq data

We received on an average 30 million 50 bases long paired end reads for each of the replicates and had two replicates for each condition. The transcript (RefSeq genes) sequences for the genome build hg38 were downloaded from UCSC table browser on Dec 10, 2016 (http://genome.ucsc.edu). We used default parameters of Kallisto (Bray et al. 2016) to build an index for the above transcript sequence and then quantified abundances of the transcripts from the paired end RNAseq fastq reads (Bray et al. 2016). The DESeq2 (Love et al. 2014) package in R was used for differential expression analysis of the quantified data obtained from Kallisto. The normalized count data from DESeq2 was used for all other downstream analysis.

We also examined expression level of genes in human cell lines with low-Myc (GEO ID = GSM1001393) & high-Myc (GEO ID = GSM1001394) condition (Loven et al. 2012). Data was analyzed using Kallisto (Bray et al. 2016) as mentioned above. The total number of estimated count (by adding estimated count of all the genes) in low and high myc condition were 85,340,683 and 106,644,300 respectively. Since number of estimated count (k-mer sequence mapping on transcripts) also depends on number of reads in a given library therefore to compare the expression of genes between the low and high myc condition the estimated count of each genes in low myc condition was multiplied by a factor 1.249 (106,644,300/85,340,683 = 1.249).

### tRF target gene prediction

The 3’UTR sequence of each annotated RefSeq genes was downloaded using UCSC Table Browser [hg38 genome build]. We decrease the noise from the lowly expressed isoform by considering only most expressed isoform of each genes in Hela cells (Nam et al. 2014) for downstream analysis. A total of 9294 sequences were examined for the complementarity of various seed sequences. Of the possible seed mers the preference was given as follows: 8mer > 7mer-m8 > 7mer-A1 > 6mer. In other words the sequences were first scanned for 8mer sites then the remaining pool was used to scan for 7mer-m8 sites and so on.

### Cumulative distribution function plot (CDF plot) of tRF target and non-target genes

Cumulative distribution function of R (ecdf) was used to compare the plot between various types of target and non-target genes in various experimental conditions. ks.test (Kolmogrov-Smirnov test), a function in R package was used to test if the plot of target genes is above the plot of non-target genes and the difference is statistically significant.

## Author contributions

C.K. and A.D. designed the experiments. C.K. is responsible for all biological experiments and P.K. is responsible for bioinformatics analysis. Initial bioinformatics analysis was done by M.K. A.M performed Ago western blots. Z.S. performed some of the biological experiments. C.K. and A.D. wrote the paper.

## Acknowledgements

We would like to thank all the members of Dutta lab for fruitful discussions. We also would like to thank for Dr. Bryan Cullen for the Dicer knock out cells and Dr. Richard Young for P493-6 cells. This work was supported by P01 CA104106 (Project 3) and R01 GM84465 to A.D. M.K. was partly supported by a DOD award (PC151085).

